# SnakeAltPromoter Facilitates Differential Alternative Promoter Analysis

**DOI:** 10.1101/2025.08.16.669128

**Authors:** Jiang Tan, Yuqing Wu, Ruteja Barve, Fuhai Li, Philip Payne, Nahyun Kong, Sheng Chih Jin, Richard Head, Yidan Sun

## Abstract

**Background:** Alternative promoter usage regulates isoform diversity in mammals, playing critical roles in development, disease, and cellular reprogramming. While Cap Analysis of Gene Expression (CAGE) enables precise transcription start site mapping, its high cost and limited coverage hinder scalability. In contrast, RNA-seq is abundant across biological contexts; several algorithms (ProActiv, Salmon, DEXSeq) infer promoter activity from these data, yet no unified, reproducible framework exists to execute, benchmark, and compare them or to scale alternative promoter analyses across large compendia.

**Results:** We developed **snakeAltPromoter**, an end-to-end Snakemake workflow that ingests raw FASTQ files, performs quality control and alignment, quantifies promoter activity using three complementary strategies (junction-based, transcript-based, and first-exon-based), classifies promoters into major, minor/alternative, and inactive categories, and conducts both differential promoter activity and usage analysis. Crucially, snakeAltPromoter integrates a systematic benchmarking module against matched CAGE profiles to reveal method-specific strengths and limitations. ProActiv showed the highest concordance with CAGE in promoter classification, activity and differential analysis, Salmon was robust at low coverage and intronless cases. Overall, the complete workflow recovered a majority of CAGE-validated active promoters and processed a 50 M-read RNA-seq sample in around 2h on a 32-core node, demonstrating both accuracy and scalability.

**Conclusions:** snakeAltPromoter is, to our knowledge, the first reproducible framework that pairs comparative method evaluation with scalable differential alternative promoter analysis. It provides concrete guidance for method selection under different experimental scenarios and enables high-throughput mining of promoter-level regulation from public RNA-seq repositories. Code and example data are freely available at https://github.com/YidanSunResearchLab/SnakeAltPromoter.git.

## Introduction

Promoters, located upstream of transcription start sites (TSSs), are pivotal in regulating transcription initiation by integrating signals from distal regulatory elements and epigenetic modifications [1–3]. In mammals, most protein-coding genes are controlled by multiple promoters, driving the expression of distinct isoforms [3–5]. Unlike post-transcriptional mechanisms such as alternative splicing, alternative promoter usage enables pre-transcriptional regulation, influencing isoform diversity [6].

Increasing evidence links alternative promoter usage to critical biological functions, underscoring its importance in development, disease, and cellular reprogramming. During organ development, alternative promoters drive organ-specific expression of isoforms, enabling precise regulation of processes like neuronal differentiation [1, 5, 7]. In disease, aberrant promoter usage is implicated in cancers. Promoters are deregulated across tissues, cancer types, and patients, affecting known cancer genes and novel candidates [8]. These diverse roles highlight the need to study alternative promoter usage, as it governs isoform diversity and regulatory complexity across biological contexts.

Cap Analysis of Gene Expression (CAGE) provides high-resolution TSS mapping, making it ideal for studying alternative promoter usage [9]. However, its high cost, technical complexity and limited coverage restrict its scalability. In contrast, RNA-seq data are abundant across biological fields, yet their potential for alternative promoter analysis remains underutilized due to the lack of standardized workflows. Moreover, computational strategies infer promoter activity from RNA-seq rely on different signals (i) splice junctions emerging from first introns (ProActiv) [8], (ii) transcript abundances (Salmon) [10], or (iii) exon-level counts (DEXSeq) [11]. No community-standard pipeline exists to easily perform these methods or guide users in selecting the optimal tool. As a result, most RNA-seq studies ignore alternative promoter usage even when their data could support it.

Here, we present snakeAltPromoter, a reproducible and extensible Snakemake workflow for differential alternative promoter analysis from RNA-seq data. The pipeline ingests raw FASTQ files, performs preprocessing and quality control, quantifies promoter activity with ProActiv, Salmon, and DEXSeq. It then classifies promoters (major vs. minor/alternative vs. inactive) and performs both differential promoter activity and differential alternative promoter usage analyses. Finally, we benchmarked each method against matched CAGE data. By coupling a systematic comparative framework with a scalable, one-command implementation, snakeAltPromoter provides actionable guidance on tool selection and unlocks promoter-level insights from the vast corpus of public RNA-seq datasets.

## Results

### Pipeline overview

snakeAltPromoter provides a comprehensive workflow for differential alternative promoter analysis. The workflow is organized into two top-level modules: (1) Genome setup (Fig. 1A) and (2) Alternative promoter analysis (Fig. 1B). The genome setup module is executed once per reference build and prepares all immutable resources, STAR [12] and Salmon [10] indices, a harmonized promoter–transcript–gene map, and auxiliary annotation files. By storing these artifacts, subsequent analyses only need to point to a directory rather than regenerate indices, which shortens turnaround time and guarantees that all downstream steps use an identical promoter definition.

**Figure 1.**
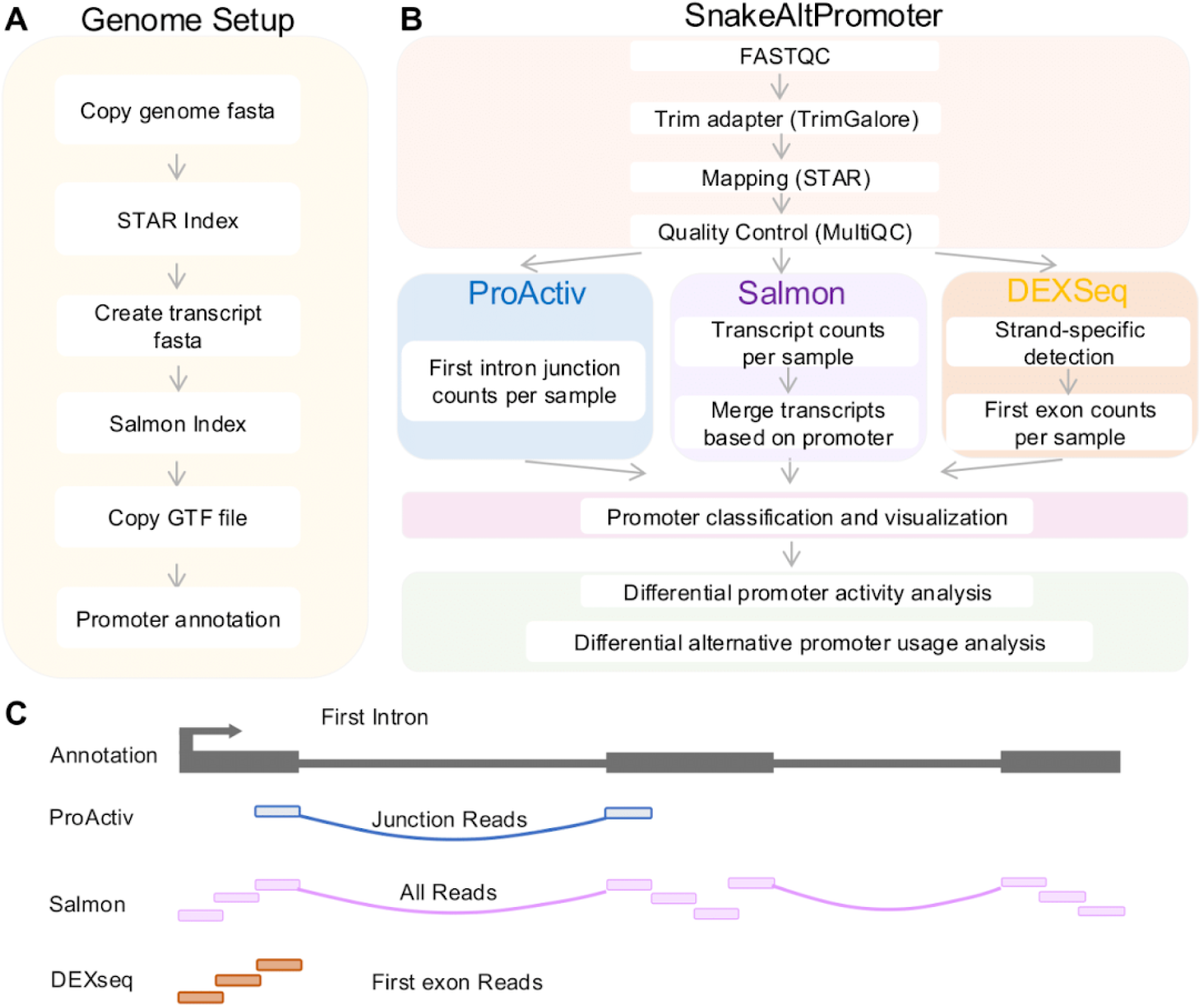
Pipeline Overview. Schematic overview of the SnakeAltPromoter pipeline. The workflow is organized into two top-level modules: Genome setup and Alternative promoter analysis. A) The genome setup module prepares reference files, including genome indices and promoter annotations. B) The analysis module processes raw FASTQ files through preprocessing, promoter quantification, classification, and differential analysis. C) Promoter counts are measured using three methods:

- ProActiv: Counts reads from first-intron junction in STAR junction files, reflecting promoter initiation.
- Salmon: Quantifies transcript abundances via quasi-mapping, with custom R scripts aggregating promoter-level counts.
- DEXSeq/featureCounts: Counts reads in first-exon bins and aggregates to promoters.

The alternative promoter analysis module processes each RNA-seq project and is itself subdivided into 4 sections (Fig. 1B). First, data preprocessing performs raw FASTQ quality assessment (FastQC) and adapter/low-quality trimming (TrimGalore), followed by STAR [12] two-pass alignment to produce coordinate-sorted BAMs and splice-junction tables. Eventually, quality results are summarized with MultiQC [13].

Second, promoter quantification is run in parallel using three complementary principles (Fig. 1C): (i) *junction-based* ProActiv [8] counts reads traversing the first intron junction, directly reflecting initiation at a given promoter and performing well when junction coverage is high; (ii) *transcript-based* Salmon [10] estimates transcript abundances via lightweight quasi-mapping and then aggregates transcripts initiating at the same promoter, offering speed and robustness for low-depth or single-end libraries; and (iii) *first-exon-based* DEXSeq [11]/featureCounts [14] measures counts of first-exon bins, capturing promoters distinguished by first exon architecture and tolerant of cases where splice junction evidence is sparse but exon boundaries are well annotated. All three methods ingest the same promoter table to ensure strict comparability.

Third, promoter classification (major, minor/alternative, inactive) is performed based on log2-tranformed promoter counts (Fig. 1B). The promoter with the highest average activity for each gene across RNA-seq samples was designated as the major promoter.

Promoters with an average activity below 0.25 were classified as inactive, while the remaining promoters with intermediate activity were categorized as minor/alternative promoters.

Lastly, differential analyses are carried out at two layers (Fig. 1B). We first identify differentially expressed promoters between conditions using DESeq2 [12]. We then test for differential alternative promoter usage, in which the relative contribution of individual promoters within a gene shifts. This second layer exposes regulatory rewiring that conventional differential expression misses and is often indicative of altered transcription factor input or chromatin accessibility.

Importantly, snakeAltPromoter can also process matched CAGE data through the STAR [15] and featureCounts steps, producing CAGE-derived promoter counts that seamlessly integrate into the workflow. This capability enables direct benchmarking of ProActiv, Salmon, and DEXSeq against gold-standard CAGE profiles, revealing method-specific strengths and limitations.

The pipeline’s scalability is a key feature. On a 32-core workstation a 50 million-read paired-end human RNA-seq sample traversed the entire snakeAltPromoter workflow in 2 h 8 min. The only memory-intensive phase was STAR’s second pass, which briefly peaked at ∼36 GB RSS; every other step, including quality control, trimming, quantification, and DESeq2, remained below 2 GB and each completed in well under a minute. Cumulative CPU time was ∼10 000 s, so the aligner effectively saturated the available cores (mean load > 130), while total I/O was modest at ∼18 GB read and ∼17 GB written, making the pipeline compute-bound rather than I/O-bound. Processing seven comparable libraries with Snakemake capped at eight concurrent jobs finished in ∼3.5 h wall time while remaining below 40 GB RAM, illustrating near-linear scaling with additional cores. This equates to an effective throughput of ∼115 million reads per hour; by simple linear extrapolation, 100 libraries of the same depth would complete in under 15 h, underscoring the pipeline’s efficiency and scalability.

SnakeAltPromoter is easy to use by utilizing snakemake’s [16] workflow management and Conda’s dependency handling [17]. With a single command line, users can execute the entire pipeline, generating results from all three quantification methods, promoter classifications, and differential analyses. Alternatively, users can select a specific quantification method (e.g., ProActiv for high-resolution studies) or modify statistical cutoffs (e.g., FDR or fold-change thresholds) via configuration parameters, offering flexibility for diverse experimental scenarios. By leveraging Conda environments, snakeAltPromoter simplifies tool installation and ensures reproducibility across systems. The modular design supports parallel processing on cluster or local platforms, making snakeAltPromoter suitable for large-scale RNA-seq datasets, such as those in public repositories like GEO or SRA. This integrated and user-friendly framework ensures reproducibility and accessibility, positioning snakeAltPromoter as a pioneering tool for promoter analysis.

### Classification of alternative promoters

We applied the pipeline to pairwise RNA-seq and CAGE data in human heart healthy and failure samples (GSE147236). Using promoters defined as regulatory regions upstream of transcription start sites (TSSs) and based on GENCODE [18] (release 46) annotations, we compiled 142,932 potential promoters (Supplementary Table S1), assuming isoforms with identical or closely spaced TSSs are regulated by the same promoter [12,13]. Focusing on promoter-level analysis reduces computational complexity compared to isoform-level analysis, enhancing robustness of inferences due to the smaller number of promoters per gene [14]. To minimize false positives, we excluded internal promoters overlapping non-first exons in certain isoforms, retaining 118,403 promoters for analysis.

Promoter activity was quantified as log2-transformed counts using ProActiv, Salmon, DEXSeq, and CAGE. Promoters were classified as *major* (highest activity per gene), *minor/alternative* (lower activity above a threshold of 0.25), or *inactive* (no detectable expression) (Fig. 2, Supplementary Table S2). Classification results varied slightly among methods. ProActiv classified 13.91% of promoters as major, 4.7% as minor/alternative, and 81.39% as inactive; Salmon yielded 25.23%, 12.14%, and 62.63%; DEXSeq produced 27.22%, 11.01%, and 61.77%; and CAGE identified 19.67%, 7.92%, and 72.42% (Fig. 2A).

**Figure 2.**
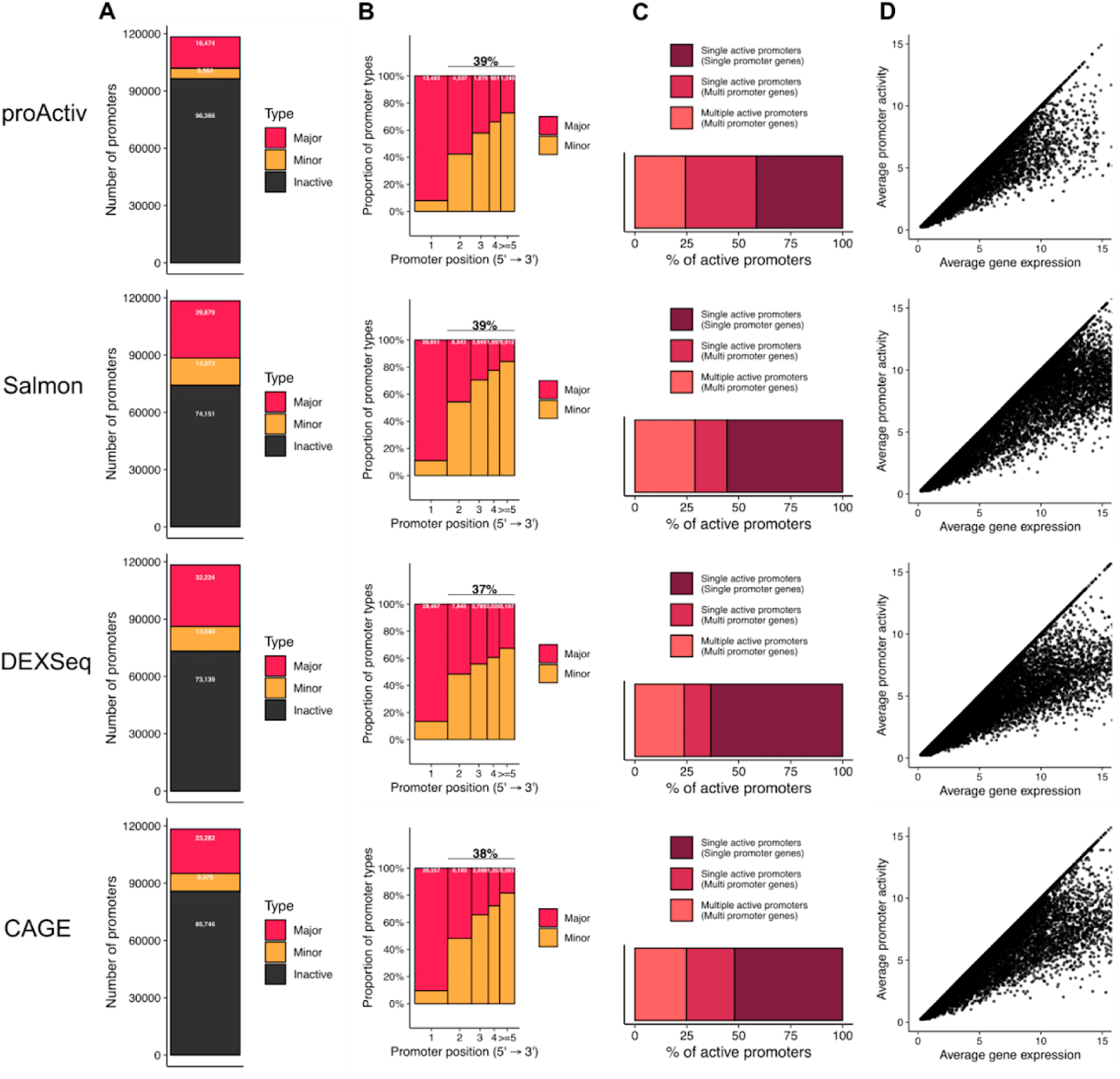
Classification of alternative promoters. (A) Classification of annotated promoters based on average promoter activity across all RNA-seq samples for each method. Promoters are categorized into three groups: major promoters, the most active promoter of each gene; minor/alternative promoters, other active promoters of the gene; and inactive promoters, with an estimated activity of <0.25. (B) Distribution of major and minor/alternative promoters across transcription start sites (TSSs), ranked from 5′ to 3′, for multi-promoter genes with at least one active promoter for each method. (C) Proportions of genes with a single promoter, a single active promoter, and multiple active promoters for each method. (D) Comparison between major promoter activity and total gene expression (sum of all promoters) for each method. A single promoter often does not fully represent the gene’s expression, as minor/alternative promoters contribute additional regulatory information.

The observed differences among methods stem from their distinct quantification strategies. DEXSeq measures first-exon bins, which can capture reads from paused polymerase or unstable short RNAs, inflating promoter counts and reducing alignment with CAGE, which primarily detects transcription initiation [11]. Salmon aggregates full-transcript abundances, diluting promoter-proximal signals with downstream exonic reads, particularly in long transcripts [10]. In contrast, ProActiv counts junction reads from the first intron, a hallmark of productive elongation following the release of promoter-proximal pausing [8]. This approach aligns closely with the mature transcript output reflected by CAGE peaks. Additionally, ProActiv’s higher inactive promoter proportion reflects its limited detection of intronless transcripts lacking introns.

Notably, the assumption that the first annotated promoter is dominant was not universally supported. ProActiv found 39% of major promoters located downstream of the first TSS, Salmon 39%, and DEXSeq 37% (Fig. 2B), consistent with prior studies [5, 8]. This finding underscores the need for empirical quantification rather than reliance on annotation-based assumptions, as dominant promoters can occur at various gene positions.

Promoter activity analysis also revealed contributions to gene expression profiles. Among expressed genes, each method identified around 25% with at least two active promoters (Fig. 2C,D). These minor/alternative promoters, often overlooked in gene-level analyses, exhibit context-dependent activity, highlighting their biological significance.

Collectively, these cross-method comparisons show that although absolute counts vary, the central conclusion—widespread alternative promoter activity—is robust. Moreover, promoter-level quantification adds an informative regulatory layer beyond genome annotations and complements conventional gene-level expression analyses.

### Benchmarking with CAGE for promoter classification

To evaluate the accuracy of ProActiv, Salmon, and DEXSeq in classifying promoters, we compared their outputs against CAGE-derived classification in human heart RNA-seq samples. We assessed the overlap in promoter classification (major and minor/alternative promoters) across methods using Venn diagrams.

For major promoters, ProActiv shared the highest fraction of its major promoters with CAGE (78.6%), reflecting its reliance on first-intron junction reads, which closely align with CAGE’s TSS-specific signals (Fig. 3A top panel). Salmon (Fig. 3A middle panel) and DEXSeq (Fig. 3A bottom panel) shared lower fractions (57.0% and 41.2%, respectively), as their transcript- and exon-based approaches include non-productive transcripts, such as paused RNAs, inflating their promoter counts. Notably, ProActiv identified the lowest number of promoters and captured only 55.6% of CAGE’s major promoters, due to its failure to detect intronless transcripts, which lack junction reads (Fig. 3A top panel). In contrast, Salmon captured 73.1% (Fig. 3A middle panel) of CAGE’s major promoters and DEXSeq captured 57% (Fig. 3A bottom panel), with Salmon’s higher fraction reflecting its ability to detect intronless genes, complementing ProActiv’s limitation. In summary, Venn diagrams showed that over 55% of promoters identified by CAGE overlapped with RNA-based methods, indicating that RNA-seq data correctly classify most major promoters (Fig. 3A).

**Figure 3.**
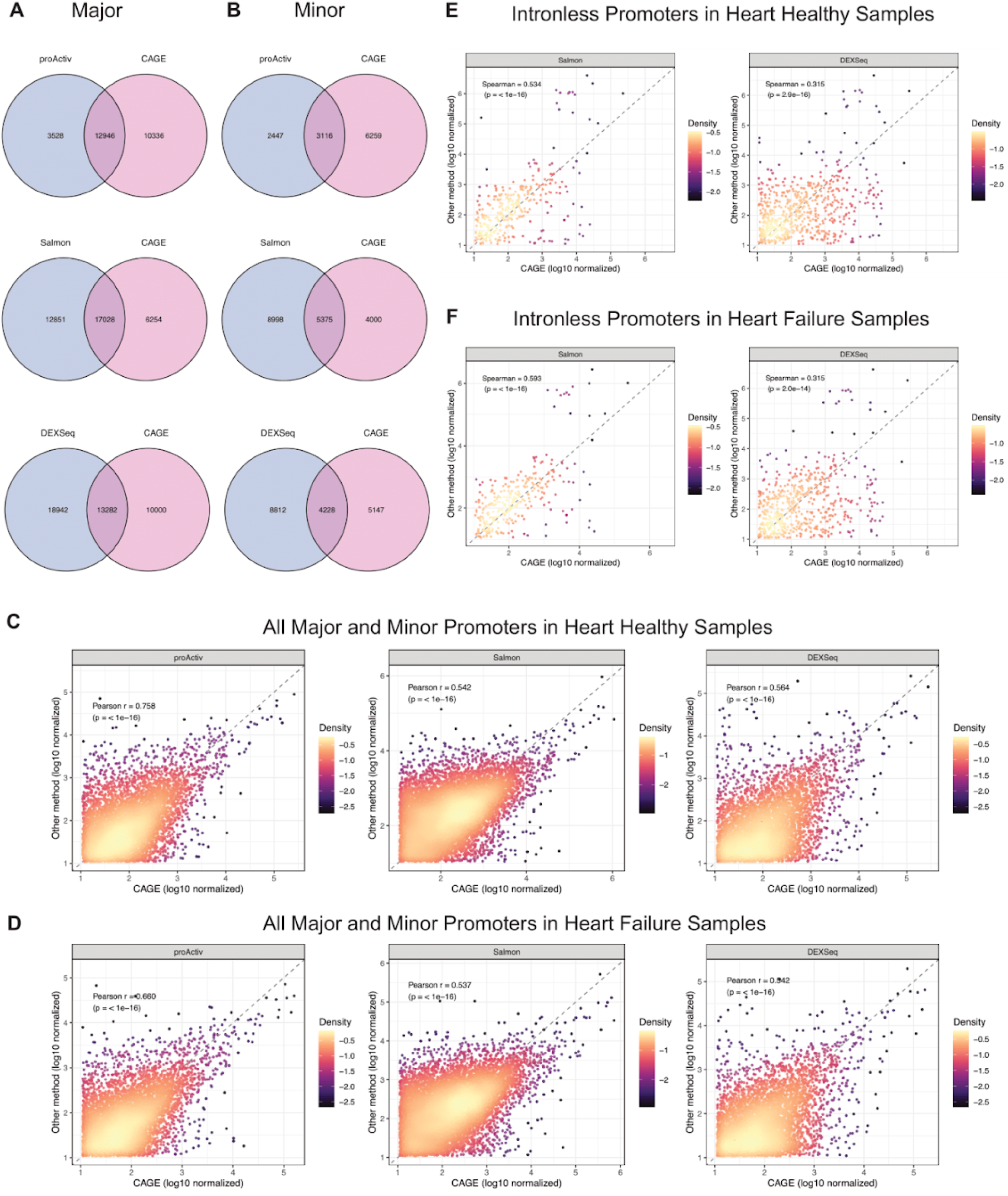
Benchmarking with CAGE for promoter classification and counts. (A) Venn diagrams showing overlap of major promoters called by CAGE and each RNA-seq method in all heart samples: ProActiv (top), Salmon (middle), and DEXSeq (bottom). (B) Same as (A), but for minor/alternative promoters. (C) Promoter-wise scatterplots comparing log10-transformed promoter counts between CAGE (x-axis) and ProActiv (left), Salmon (middle), and DEXSeq (right) for all promoters in healthy heart samples; each plot is annotated with the Pearson correlation coefficient (PCC) and associated *P*-value. (D) Same as (C), but for heart failure samples. (E) Scatterplots of intronless promoter counts in healthy heart samples, comparing CAGE versus Salmon (left) and CAGE versus DEXSeq (right); plots are annotated with PCC and *P*-value. (F) Same as (E), but for heart failure samples.

For minor/alternative promoters (Fig. 3B), ProActiv again shared the highest fraction with CAGE (56.0%), while Salmon and DEXSeq had lower fractions (37.4% and 32.4%, respectively), consistent with the trend for major promoters. Similarly, when assessing the fraction of CAGE-defined minor/alternative promoters captured, Salmon identified the highest fraction (57.3%, Fig. 3B middle panel), followed by DEXSeq (45.1%, Fig. 3B bottom panel), with ProActiv (33.2%, Fig. 3B top panel) capturing the lowest due to its limitation with intronless genes. Notably, the overlap in minor/alternative promoters is lower compared to major promoters, indicating method-specific sensitivities to low-coverage or complex 5’ regions.

Overall, identification of both major and minor promoters from RNA-seq was largely consistent with CAGE. Detection of minor/alternative promoters was weaker than that of major promoters, likely due to their lower expression. ProActiv shares the highest fraction of its promoters with CAGE for both promoter types, likely due to its exclusion of paused or unstable transcripts. However, Salmon and DEXSeq complement ProActiv by detecting intronless genes, with Salmon capturing the highest fraction of CAGE promoters. These findings validate RNA-seq’s ability to recover CAGE-defined promoter classes, with ProActiv optimal for matching CAGE signals and Salmon preferred for capturing intronless promoters, guiding method selection based on experimental goals.

### Benchmarking with CAGE for promoter counts

To assess how closely RNA-seq-derived promoter counts align with CAGE, we compared the outputs of ProActiv, Salmon, and DEXSeq against CAGE-derived counts in human heart RNA-seq samples (Supplementary Table S3). For each method, we generated scatterplots of promoter-wise counts against CAGE, annotated with Pearson correlation coefficients (PCC) and associated P-values (Fig. 3C,D). Across all major and minor promoters in healthy heart samples (Fig. 3C), ProActiv achieved the highest median PCC (0.758), followed by DEXSeq (0.564) and Salmon (0.542), all with P < 1e-16. Heart failure samples yielded similar results (Fig. 3D). These data confirm that ProActiv exhibits the strongest concordance with CAGE, in line with its superior performance in promoter classification.

A key limitation of ProActiv is its inability to quantify intronless promoters, which lack first-intron junctions and number over 20,000 in the human genome. To address this, we analyzed intronless promoters using only Salmon and DEXSeq, assessing their counts against CAGE with Spearman correlation to account for non-linear relationships in these data (Fig. 3E,F). In healthy heart samples (Fig. 3E), Salmon’s counts correlated more closely with CAGE (correlation=0.534) than DEXSeq’s (correlation=0.315). Comparable results were observed in heart failure samples (correlation = 0.593 for Salmon vs. 0.315 for DEXSeq) (Fig. 3F), indicating that DEXSeq’s exon-based counts are more susceptible to paused or non-productive transcription signals, while Salmon’s transcript-based approach better captures intronless promoter activity.

In summary, ProActiv consistently outperforms Salmon and DEXSeq in aligning RNA-seq-derived promoter counts with CAGE across all promoters, driven by its junction-based approach that closely tracks productive transcription. However, for intronless promoters, which ProActiv cannot detect, Salmon provides more reliable quantification than DEXSeq, whose inclusion of paused transcripts reduces accuracy. These findings highlight ProActiv’s strength for most promoter analyses and Salmon’s utility for intronless cases, guiding tool selection within the snakeAltPromoter pipeline.

### Benchmarking with CAGE for differential alternative promoter analysis

To benchmark each method’s ability to detect differential promoter activity, we first contrasted healthy and failing human heart samples. Differential promoter analysis was performed using DESeq2 on promoter count matrices derived from CAGE, ProActiv, Salmon, and DEXSeq (Supplementary Table S4). As expected, the most significant changes were losses of promoter activity in heart failure in the CAGE analysis (Fig. 4A). Pathway analysis of downregulated promoters in failing versus healthy hearts revealed coordinated repression of pathways central to cardiac structure and function. Signals converged on cardiomyopathy programs with prominent involvement of sarcomere and myofibril organization, consistent with loss of contractile integrity. The dominant themes included cardiomyopathy, sarcomeres/myofibrils, signal conduction, and axon/myelin maintenance. A hypertrophic cardiomyopathy module was especially prominent, with leading genes PDE4A, TNNT2, AKAP13, CAV1, SLC4A3, ADRB1, FHOD3, ASB15, EIF4BP1, SMYD1, TMP1, ASB2, ACHYL1, and PKIA enriched, underscoring that promoter-level repression in heart failure converges on canonical pathways of adverse cardiac remodeling (Fig. 4B).

**Figure 4.**
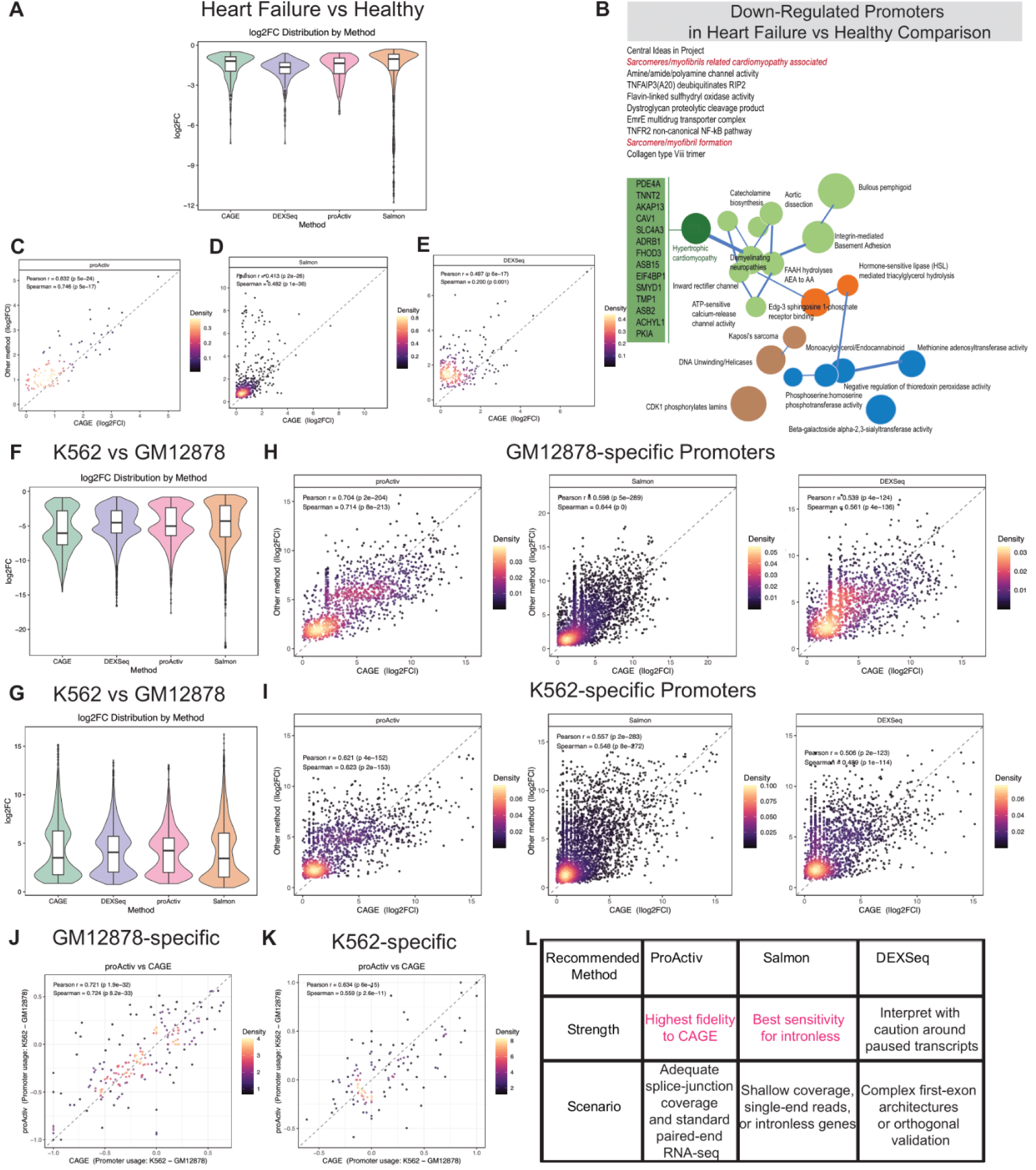
Benchmarking with CAGE for differential alternative promoter analysis. (A) Violin plot showing log₂ fold-changes (log2FC) of down-regulated promoters between healthy and failing heart samples as detected by ProActiv, Salmon, DEXSeq and CAGE (FDR < 0.05). (B) Pathway map of down-regulated promoters in failing heart samples compared to healthy controls using the CompBIo (v2.9) platform. The resulting map illustrates key pathways and processes associated with heart failure. Prominent themes include hypertrophic cardiomyopathy, sarcomere/myofibril organization, signal conduction, and axon/myelin maintenance. Pathways are color-coded as follows: signal conduction (green), metabolic processes (blue), DNA and nuclear processes (brown), and lipid metabolism (orange). The hypertrophic cardiomyopathy theme (dark green) is highlighted, along with its top enriched genes. (C,D,E) Promoter-wise scatterplots comparing log2FC of down-regulated promoters (heart failure vs. healthy) between CAGE (x-axis) and ProActiv (B), DEXSeq (C) or Salmon (D)(y-axis); each panel is annotated with the Pearson correlation coefficient (PCC) and its *P*-value. (F) Violin plot showing log2FC of GM12878-specific promoters, which are down-regulated promoters (FDR < 0.05) identified by ProActiv, Salmon, DEXSeq and CAGE in the K562 vs. GM12878 comparison. (G) Violin plot showing log2FC of K562-specific promoters, which are up-regulated promoters (FDR < 0.05) detected by each method and CAGE in the K562 vs. GM12878 comparison. (H) Scatterplots of log2FC for GM12878-specific promoters (down-regulated in K562 vs. GM12878): CAGE (x-axis) versus ProActiv (left), Salmon (middle), and DEXSeq (right) (y-axis), with PCC and *P*-value indicated. (I) Scatterplots of log2FC for K562-specific promoters (up-regulated in K562 vs. GM12878): CAGE (x-axis) versus ProActiv (left), Salmon (middle), and DEXSeq (right) (y-axis), annotated with PCC and *P*-value. (J) Scatterplot comparing GM12878-specific promoter usage shifts in each gene between GM12878 and K562 as measured by CAGE (x-axis) and ProActiv (y-axis); the PCC and *P*-value are shown. (K) Scatterplot comparing K562-specific promoter usage shifts in each gene between GM12878 and K562 as measured by CAGE (x-axis) and ProActiv (y-axis); the PCC and *P*-value are shown. (L) Summary table outlining recommended use cases for each RNA-seq–based quantification method.s

We then compared log2 fold change (log2FC) between RNA-seq–based methods and CAGE. ProActiv showed the highest correlation with CAGE (PCC = 0.832), followed by DEXSeq (PCC = 0.497) and Salmon (PCC = 0.413), all with P < 1e-16 (Fig. 4C,D,E). These results demonstrate that junction-based quantification not only aligns with CAGE in static promoter classification but also faithfully captures dynamic changes in promoter activity during disease.

To evaluate method performance in a different biological context, we next obtained matched RNA-seq and CAGE data for the GM12878 and K562 cell lines from ENCODE (GSE33480 and GSE34448). We compared K562 to GM12878 and identified up-regulated promoters as K562-specific promoters and down-regulated genes as GM12878-specific promoters (Fig. 4F,G). Scatterplots of log2FC values for these promoters were generated against CAGE, with PCCs calculated to assess agreement. For GM12878-specific promoters, ProActiv showed the strongest concordance with CAGE (Pearson r = 0.704), followed by Salmon (r = 0.598) and DEXSeq (r = 0.539); all correlations were highly significant (P < 1 × 10⁻¹⁶; Fig. 4H). The same pattern emerged in K562-specific promoters: ProActiv again led (r = 0.621), with Salmon (r = 0.557) and DEXSeq (r = 0.506) close behind (Fig. 4I). These results confirm that the relative performance of the three methods is consistent across cell types. ProActiv’s strong alignment with CAGE confirms the robustness of ProActiv’s junction-based approach in capturing dynamic promoter activity across diverse sample types.

To assess RNA-seq’s ability to detect alternative promoter usage, we focused on ProActiv due to its strong correlation with CAGE counts. Promoter usage was calculated for each cell type as the ratio of individual promoter counts to total gene expression counts. Usage shifts were determined by subtracting GM12878 promoter usage from K562 usage for each differentially expressed promoter identified previously. Promoter usage shifts more than 0.1 were kept for further analysis. Scatterplots of the promoter usage shift values were generated, comparing ProActiv against CAGE, with Pearson correlation coefficients (PCCs) calculated to evaluate agreement. For GM12878-specific (Fig. 4J) and K562-specific promoters (Fig. 4K), ProActiv showed high concordance with CAGE (PCC = 0.721 and 0.634, respectively, P < 1e-16). These findings validate ProActiv’s reliability in detecting context-specific promoter usage changes from RNA-seq data.

Overall, ProActiv consistently outperforms Salmon and DEXSeq in aligning differential promoter activity and alternative promoter usage with CAGE across heart and cell line datasets, driven by its ability to track productive transcription. Consequently, for large-scale alternative promoter analyses where CAGE is unavailable, ProActiv provides a robust surrogate for detecting biologically meaningful promoter switches.

### Summary

To our knowledge, snakeAltPromoter is the first reproducible framework that couples comparative method evaluation with scalable differential alternative promoter analysis, unifying ProActiv, Salmon, and DEXSeq in a single Snakemake workflow. What distinguishes snakeAltPromoter is that it (i) feeds all three methods the same harmonised promoter table, eliminating coordinate mismatches; (ii) benchmarks their outputs against optional CAGE data in the same run, giving users an on-the-fly accuracy report; (iii) scales from a laptop to an HPC node with near-linear speed-ups; and (iv) stores every step, parameter and software version under Snakemake and Conda for full provenance. All together, snakeAltPromoter turns complex promoter-level analysis into a single, reproducible command.

In addition, snakeAltPromoter provides actionable guidance for method choice. For datasets with ample splice-junction coverage and standard paired-end reads, ProActiv is the first-choice quantifier; when sequencing is shallow, single-end, or enriched for intron-poor genes, Salmon provides the closest match to CAGE; and DEXSeq, while niche, is invaluable for questions centred on first-exon architecture (Fig. 4L). Users can therefore default to ProActiv, fall back to Salmon for intronless focus, and deploy DEXSeq for exon-architecture checks (Fig. 4L)—confident that whichever path they choose, the surrounding QC, classification, differential testing and documentation are handled automatically by snakeAltPromoter.

## Methods

### Pipeline architecture

snakeAltPromoter is implemented in Snakemake [16], providing a readable, scalable workflow that runs on local workstations, HPC clusters, and cloud platforms. All third-party software is distributed through Conda/Bioconda [17], eliminating manual dependency management and the need for administrator privileges.

### Genome setup

For GRCh38 (GENCODE [18] v46) we downloaded the primary FASTA and comprehensive GTF. The FASTA was indexed for STAR [15] and Salmon [10]. Promoter coordinates were generated with preparePromoterAnnotation from proActiv [8], merging overlapping first exons and keeping the 5′-most TSS. Internal promoters that overlap downstream exons of other transcripts were excluded. The resulting promoter-to-transcript-to-gene map is reused in every downstream step, ensuring consistent promoter definitions.

### Data Preprocessing

Raw FASTQ files undergo quality assessment with FastQC (https://www.bioinformatics.babraham.ac.uk/projects/fastqc/), followed by adapter and low-quality base trimming (Q < 20) using TrimGalore (https://github.com/FelixKrueger/TrimGalore). Trimmed reads are aligned to the reference genome with STAR [15] in two-pass mode, producing coordinate-sorted BAMs and splice-junction tables.

### Promoter Counts Quantification

Promoter counts are measured using three methods:

- ProActiv [8]: Counts reads from first-intron junction in STAR junction files, reflecting promoter initiation.
- Salmon [10] : Quantifies transcript abundances via quasi-mapping, with custom R scripts aggregating promoter-level counts.
- DEXSeq/featureCounts [11]: Counts reads in first-exon bins and aggregates to promoters.

All three methods use the same promoter annotation, guaranteeing strict comparability. Only uniquely mapped reads are used for counting in each method.

### Definition of major, minor/alternative, inactive promoters

We categorized the promoters into three distinct groups (major, minor/alternative, and inactive) based on their log2-transformed counts using getAlternativePromoters from proActiv [8]. The promoter with the highest average activity for each gene across all RNA-seq samples was designated as the major promoter. Promoters with an average activity below 0.25 were classified as inactive, while the remaining promoters with intermediate activity were categorized as minor/alternative promoters.

### Differential Promoter activity analysis

For each quantification method, we applied DESeq2 [12] to identify differentially expressed promoters using a FDR cutoff of 0.05.

### Differential alternative Promoter usage analysis

For each quantification method, promoter usage was calculated as the ratio of individual promoter counts to total gene expression counts. Usage shifts were determined by subtracting promoter usage between 2 conditions for each differentially expressed promoter identified previously. We also apply getAlternativePromoters from Proactiv to calculate the adjusted P-value.

### CAGE data processing

Raw CAGE FASTQ files are subjected to the identical preprocessing and alignment workflow used for RNA-seq. Briefly, FastQC evaluates read quality, TrimGalore removes adapters and low-quality bases, and STAR [15] is run in two-pass mode to align reads to the reference genome. Promoter counts are then obtained by running featureCounts against the promoter annotation used for RNA-seq quantification. The resulting promoter-by-sample count matrix is processed through the same promoter classification and differential analysis modules: promoters are labeled as major, minor/alternative, or inactive based on log2-transformed counts, and DESeq2 [12] is used to identify differentially active promoters (FDR < 0.05). promoter usage was calculated and usage shifts were determined using the same method introduced previously. This consistent treatment ensures that CAGE data integrate seamlessly into all downstream promoter classification and differential promoter analysis comparisons.

### CompBio pathway analysis

Promoters that were significantly downregulated in failing versus healthy hearts (DESeq2, FDR < 0.05, log₂FC < 0) were mapped to gene symbols and supplied to CompBio v2.9 (https://gtac-compbio.wustl.edu/legacy/resource/). CompBio builds a literature-derived knowledge base from PubMed abstracts and full-text articles using natural language processing and conditional probability, then evaluates the input gene list against this knowledge base to identify statistically enriched biological concepts.

Concepts are aggregated into context-aware biological themes that represent pathways and processes; theme size and rank reflect the absolute enrichment between concepts and the genes assigned to them, and proximity between themes reflects shared membership. We used default settings and retained themes that met CompBio’s significance criteria (normalized enrichment score > 1.2 and empirical p value < 0.1), as computed by comparing each theme’s absolute enrichment to themes of the same rank in thousands of randomized gene lists of matched size. Theme annotations were assigned by CompBio in a fully contextualized manner, considering both the concepts within a theme and the relationships to neighboring themes. For reporting, we exported the ranked theme table and the knowledge map and summarized leading genes within selected themes that were most relevant to heart failure biology.

### Benchmarking against CAGE

To evaluate the accuracy of ProActiv, Salmon, and DEXSeq in promoter analysis, we compared their outputs against pairwise CAGE-derived data using human heart samples. We performed three types of comparisons: promoter classification, promoter count estimation, and differential promoter activity.

### Promoter classification comparison

We evaluated the overlap of major and minor/alternative promoters identified by each method with CAGE using Venndiagram [19]. We calculated the fraction of each method’s promoters shared with CAGE and the fraction of CAGE-defined promoters captured by each method to assess classification accuracy.

### Promoter counts comparison

We generated scatterplots of promoter-wise counts for each method against CAGE-derived counts using ggplot2 [20]. To reduce noise, we normalized all the counts using DESeq2 [12] and filtered out promoters with normalized counts below 10. We calculated Pearson correlation coefficients (PCC) and associated P-values to assess the correlation between RNA-seq and CAGE counts using R basic functions.

For intronless promoters, which lack first-intron junctions and cannot be quantified by ProActiv, we generated scatterplots of promoter-wise counts for only Salmon and DEXSeq against CAGE-derived counts using ggplot2 [20]. We calculated Spearman correlation coefficients and associated P-values to account for non-linear relationships in count data.

### Differential alternative promoter analysis comparison

For differential promoter activity, we computed absolute log2 fold changes (log2FC) between healthy and heart failure samples for each method using DESeq2 [12] (FDR < 0.05) and generated scatterplots [20] against CAGE’s absolute log2FC values. We calculated PCCs and P-values to evaluate the concordance of differential promoter activity between RNA-seq and CAGE.

To identify cell type-specific promoters, we obtained RNA-seq and CAGE datasets for GM12878 and K562 cell lines from ENCODE. We identified promoters specific to each cell type by comparing their expression between K562 and GM12878 and using DESeq2 (FDR < 0.05). Promoters were classified as K562-specific if log2FC > 0 or GM12878-specific if log2FC < 0 in K562 versus GM12878 comparisons. Scatterplots [20] of log2FC values for these cell type-specific promoters were generated, comparing ProActiv, Salmon, and DEXSeq against CAGE, with Pearson correlation coefficients (PCCs) and P-values calculated to assess concordance in detecting cell type-specific promoter activity.

To detect condition-dependent promoter usage shifts, we first computed promoter usage for each cell type as the ratio of individual promoter counts to total gene expression counts. We then calculated the usage shift by subtracting the GM12878 promoter usage from that of K562 for each promoter. The promoter usage shift less than 0.1 was removed. To evaluate concordance with CAGE, we performed scatterplots [20] the promoter usage shifts identified by ProActiv against those from CAGE and computed Pearson correlation coefficients with associated *P*-values.

## Discussion

snakeAltPromoter addresses a critical gap in transcriptomics by providing the first systematic pipeline for alternative promoter analysis using RNA-seq data. Unlike existing tools, which focus on gene expression or splicing, our pipeline targets promoter-level regulation, revealing isoform diversity overlooked in standard analyses. By integrating ProActiv, Salmon, and DEXSeq, snakeAltPromoter enables comparative evaluation to CAGE data in promoter classification, quantification and differential analysis. This benchmarking is a key contribution, offering researchers clear guidance on tool selection: ProActiv for high-resolution studies of intron-containing promoters, Salmon for intronless promoters and shallow sequencing analyses. While CAGE remains the gold standard for TSS mapping, snakeAltPromoter’s ability to leverage RNA-seq data democratizes promoter analysis, reducing reliance on costly technologies. The pipeline’s scalability and reproducibility, enabled by Snakemake and Conda, make it suitable for large-scale RNA-seq datasets, a critical feature given the abundance of such data in public repositories.

### Expansion to Long-Read Sequencing Data

A promising future direction for snakeAltPromoter is its adaptation to long-read sequencing data, such as those generated by PacBio [21] or Oxford [22] Nanopore platforms. Long-read sequencing offers direct resolution of full-length transcripts, overcoming limitations of short-read RNA-seq, such as ambiguous read mapping at exon junctions or in regions with complex splicing [21, 22]. ProActiv’s junction-based approach [8], which relies on first-intron junction reads, can be extended to long-read data by leveraging full-length transcript alignments to precisely identify TSSs and their associated junctions. This adaptation is likely to enhance accuracy, as long reads provide unambiguous evidence of isoforms, potentially improving detection of low-abundance minor/alternative promoters that are challenging with short reads [21, 22]. Integrating long-read support into snakeAltPromoter would involve updating the preprocessing module to handle long-read alignment tools (e.g., minimap2 [23]) and modifying ProActiv’s junction detection to process full-transcript data, offering a robust framework for studying promoter dynamics in complex transcriptomes.

### Integration with Epigenomic Data

Integrating epigenomic data—such as ATAC-seq [24] or ChIP-seq for promoter-associated histone marks (H3K4me3, H3K27ac)—could further enhance snakeAltPromoter’s capacity to characterise promoter activity. These data provide orthogonal evidence of active TSSs because open chromatin and activating histone modifications reliably mark promoter regions [24]. Incorporating such datasets would allow the pipeline to validate RNA-seq-derived promoter calls, a benefit that is especially important for minor or low-abundance promoters where RNA-seq–only methods show reduced concordance with CAGE. For instance, overlaying ProActiv’s junction-based counts with ATAC-seq peak intensity could confirm promoter activity in low-expression contexts and thereby improve classification accuracy. Implementation would require an additional module that aligns epigenomic reads, calls peaks, and intersects those peaks with the existing promoter atlas; machine-learning models could further combine multi-omics data to predict promoter status. Finally, extending the ATAC-seq branch to include digital footprinting (e.g. TOBIAS [25] or HINT-ATAC [26]) would identify transcription-factor binding events within promoter peaks, directly linking specific TFs to promoter usage shifts uncovered by snakeAltPromoter and providing mechanistic insight into tissue- or disease-specific regulatory programmes.

### Conclusions

snakeAltPromoter fills a gap left by existing promoter-quantification tools by wrapping the entire analytical chain—pre-processing, three orthogonal quantification strategies, promoter classification, differential expression, and usage testing—into one reproducible Snakemake workflow. Its unified promoter table guarantees that ProActiv, Salmon, and DEXSeq are benchmarked on an identical coordinate system, eliminating the inconsistencies that have hampered cross-tool comparisons. The built-in CAGE module further provides an objective yard-stick, showing that ProActiv is most accurate for junction-rich, intron-containing genes, whereas Salmon maintains superior recall for intron-poor loci; DEXSeq remains ideal when promoters have complex exon architecture.

Beyond accuracy, snakeAltPromoter delivers practical advantages demanded by the community. A single command launches a fully containerised run that finishes a 50 M-read sample in just about two hours on a 32-core workstation while using < 40 GB RAM—a throughput of ∼115 M reads h⁻¹ that scales nearly linearly across additional cores and projects. Command-line parameters let users swap quantifiers or tighten FDR thresholds without touching the workflow logic, making the pipeline equally suitable for exploratory screens and high-resolution studies. Freely available and scalable, snakeAltPromoter is poised to advance transcriptomics research by leveraging the wealth of RNA-seq data. By converting heterogeneous RNA-seq datasets into promoter-centric insights, it enables researchers to tap into a largely unexploited regulatory layer.

### Data Availability

The snakemake pipeline and example data are available at https://github.com/YidanSunResearchLab/SnakeAltPromoter.git.

All the RNA-seq and CAGE datasets used in the paper are from GEO with the accession codes GSE147236, GSE33480 and GSE34448.

## Abbreviations

RNA-seq: RNA sequencing
CAGE: Cap Analysis of Gene Expression
FDR: false discovery rate
PCC: Pearson correlation coefficient

## Acknowledgements

We thank Dr. Jared Lalmansingh for the technical support. We thank Dr. Jeffrey Milbrandt and Dr. Po-Yin Yen for insightful discussions and continuous support of this project. We thank all members of our department and institutes for fostering a collaborative and supportive research environment.

## Authors’ contributions

YS, JT and RH conceived the study, designed the snakeAltPromoter pipeline. JT and YW developed Snakemake rules and implemented benchmarking modules for promoter quantification and classification. RB performed pathway enrichment analysis for differentially expressed promoters. FL assisted in interpreting promoter activity and differential expression data, providing biological context for promoter analyses. PY assisted in computational strategies for alternative promoter usage analysis, enhancing pipeline analytics. NK and SCJ provided expertise in heart failure and promoter biology, shaping data interpretation. RH supervised the study, and directed strategic integration of methods. JT, YS and RH drafted the manuscript with input from all authors. All authors participated in discussions to refine pipeline design and results, and approved the final manuscript.

## Ethics declarations

### Ethics approval and consent to participate

Not applicable.

### Consent for publication

Not applicable.

## Supplementary Files

Supplementary Table S1. A comprehensive list of promoter coordinates.

Supplementary Table S2. Promoter classifications of major and minor using Proactiv, Salmon, DEXseq and CAGE.

Supplementary Table S3. Promoter counts across samples measured by Proactiv, Salmon, DEXseq and CAGE.

Supplementary Table S4. Differential promoter activity analysis between heart healthy and failure samples measured by Proactiv, Salmon, DEXseq and CAGE.

